# LINGUINE: a phylogeny-aware, orthogroup-based framework for robust ancestral linkage group inference

**DOI:** 10.64898/2026.05.26.727789

**Authors:** Carlos Vargas-Chávez, Rosa Fernández

## Abstract

Reconstructing the chromosomal architecture of ancestral genomes is a central challenge in comparative genomics, yet existing methods for ancestral linkage group (ALG) inference struggle in clades with rapid genome evolution. Current approaches rely predominantly on single-copy orthologous genes to anchor linkage groups between species, limiting marker density and lacking mechanisms to resolve paralogy, which leads to systematic inflation of inferred ALG counts and propagates structural artifacts in downstream analyses. Here we present LINGUINE (LINkage GroUps INfErence), a phylogeny-aware pipeline for robust ALG reconstruction addressing these limitations. LINGUINE uses orthogroups to maximise marker density, employs a Hidden Markov Model to delineate syntenic blocks while accommodating gene loss, rearrangements, and assembly fragmentation, and incorporates an explicit paralogy resolution step prior to ancestral reconstruction. Ancestral states are inferred through iterative post-order traversal of the species tree, progressively integrating information from all descendants. Using GARLIC (Genome reARrangement simuLator for Inferring Chromosomal landscapes), we characterise the limits of synteny signal detectability and define the conditions under which ALG reconstruction remains reliable. Benchmarking on nematodes demonstrates concordant performance with existing tools while incorporating a substantially larger fraction of the gene repertoire. Applied to clitellate annelids, a clade with highly rearranged genomes, LINGUINE recovers biologically coherent ancestral architectures that existing methods fail to resolve. These results establish LINGUINE as a flexible framework for ancestral genome reconstruction in clades where traditional approaches lose resolution. LINGUINE and GARLIC are open source and available in GitHub (https://github.com/MetazoaPhylogenomicsLab/Vargas-Chavez-Fernandez_2026_LINGUINE, and https://github.com/MetazoaPhylogenomicsLab/Vargas-Chavez-Fernandez_2026_GARLIC).

## Introduction

The organisation of genes into chromosomes is not static over evolutionary time. Chromosomal fusions, fissions, translocations, gene loss and whole-genome duplications (WGDs) continuously remodel genome architecture, and reconstructing this history is fundamental to understanding gene family evolution, chromosomal architectural dynamics, and the tempo of genomic change across lineages (Nakatani and McLysaght 2017). Ancestral linkage group (ALG) inference, the reconstruction of the chromosomal units present in an extinct ancestor based on conserved gene co-localisation across descendant lineages, provides a direct window into these processes, enabling researchers to track large-scale synteny conservation and disruption across divergent taxa. Despite growing interest in macrosynteny as a comparative framework, robust ALG inference remains a significant methodological challenge, particularly in clades characterised by rapid genome evolution, extensive chromosomal rearrangements, or polyploidisation events, precisely the biological contexts where such inference would be most informative.

The foundations of ALG inference were laid by Putnam et al. (2008), who identified 17 conserved chordate linkage groups shared between amphioxus and vertebrates, establishing the core principle that syntenically equivalent chromosomal units shared across distantly related lineages must reflect chromosomes present in their common ancestor (Putnam et al. 2008). This work demonstrated that macrosynteny, the chromosome-scale conservation of ortholog distributions, retains a detectable historical signal even when fine-scale gene order has been entirely disrupted, providing the conceptual basis for all subsequent ALG reconstruction efforts. This framework was substantially advanced by Nakatani and McLysaght (2017), who formalised macrosynteny inference into a rigorous probabilistic model based on variational Bayes inference (conceptually analogous to latent Dirichlet allocation in document analysis) and applied it to reconstruct the pre-teleost WGD ancestral genome (Nakatani and McLysaght 2017). Notably, their approach demonstrated effectiveness in precisely the situation where gene-order-based algorithms fail; ancient WGDs characterised by high rates of rearrangement and extensive gene loss, while also introducing explicit quantification of reconstruction confidence, a key advance over previous ad hoc methods. This probabilistic framework was subsequently extended by Nakatani et al. (2021), who applied it to reconstruct the proto-vertebrate, proto-cyclostome, and proto-gnathostome genomes using chromosome-scale assemblies of lamprey and elephant shark (Nakatani et al. 2021). Importantly, validation of the proto-vertebrate reconstruction against invertebrate genomes, including scallop (*Pecten maximus*) and a placozoan (*Trichoplax adhaerens)*, revealed strong macrosynteny conservation, demonstrating that the reconstructed ancestral vertebrate chromosomes had already existed as distinct units deep in metazoan evolution, predating the vertebrate lineage itself. Building on these foundations, Simakov et al. (2022) extended the inference of ancestral chromosomal units to two more invertebrate phyla (sponges and cnidarians), defining ancestral bilaterian and metazoan linkage groups traceable across cnidarians, bilaterians, sponges and mollusks (Simakov et al. 2022). Together, these studies established macrosynteny as a powerful lens through which deep animal genome evolution can be studied, independent of gene order conservation.

Despite these advances, the probabilistic frameworks of Nakatani and McLysaght (2017) and Nakatani et al. (2021) and the comparative approach of Simakov et al. (2022) share important limitations that constrain their applicability in complex genomic contexts. The macrosynteny model of Nakatani and colleagues was designed specifically around WGD events in a vertebrate context and requires a non-WGD outgroup species as an anchor, limiting its generalisability to the full diversity of metazoan genome architectures. Simakov et al. (2022), while extending ALG reconstruction beyond vertebrates to include representative invertebrate lineages, similarly relies on a restricted set of species and on single-copy orthologous genes as markers of syntenic conservation, an assumption that becomes increasingly problematic in lineages with elevated gene loss, large multigene families, or rapid sequence divergence, where the number of usable single-copy markers collapses to a fraction of the gene space. Neither framework is furthermore designed to handle the marker density problem that arises in clades with highly rearranged genomes: when the number of interchromosomal rearrangements between compared lineages is high, pairwise signal becomes ambiguous, single-copy marker sets become sparse, and the clean one-to-one correspondences underpinning reconstruction algorithms break down. Addressing this marker density problem is therefore a prerequisite for any method intended to operate robustly across phylogenetically and genomically diverse datasets.

Existing tools for ALG inference carry additional structural limitations that compound these challenges. Odp (Oxford Dot Plots), introduced by Schultz et al. (2023) as a formalisation of the pairwise synteny methodology underlying the Simakov et al. study, performs synteny assessment between extant species and was notably applied to explore the longstanding question of ctenophore phylogenetic placement through macrosyntenic characters (Schultz et al. 2023). While effective for such pairwise and small-scale multi-species comparisons, odp is limited to only a handful of species (three to five) due to the stringent ortholog selection process. Critically, odp does not reconstruct ancestral nodes, comparing extant chromosomal configurations directly and making it unsuitable for inferring the chromosomal complement of common ancestors in multi-species phylogenies. A more recent tool to infer ALGs, Syngraph (Mackintosh et al. 2023), addresses this by enabling node-by-node ancestral reconstruction across a phylogeny using a fusion–fission algorithm, and represents a significant advance in the field. However, Syngraph’s reconstruction at each internal node relies on a fixed three-species window (the two daughter lineages and a single outgroup), making ancestral state inference highly sensitive to the genomic properties of those three taxa. When any one of them carries missing data, assembly fragmentation, or an atypical chromosomal configuration resulting from lineage-specific rearrangements, the inferred ancestral state can be substantially distorted, with no mechanism to draw on the broader phylogenetic context. On the other hand, macrosyntR (Anon) takes a complementary approach by using ortholog tables and BED-formatted genome annotations to compute significantly conserved chromosome–chromosome associations via Fisher’s exact tests, and to visualise ortholog distributions on automatically ordered Oxford grids. A network-based greedy community detection algorithm is used to cluster chromosomes that likely derive from the same ancestral linkage group, improving interpretability even when one genome is fragmented. However, macrosyntR operates on pairs of extant genomes at a time, does not implement an explicit phylogeny-wide ancestral reconstruction scheme, and, like odp, is designed around single-copy orthologous markers. As a result, all three tools struggle in clades with extensive genome rearrangement, pervasive paralogy, or ancient WGDs, where marker density and paralog resolution become limiting.

In this context, we present LINGUINE (LINkage GroUps INfErence)(Fig. 1), a phylogeny-aware computational pipeline that addresses these limitations through three key innovations. First, LINGUINE uses orthogroups (OGs) or hierarchical orthogroups (HOGs) rather than single-copy genes as the unit of syntenic inference, substantially increasing marker density and retaining signal in divergent or fast-evolving lineages where BUSCO-based approaches lose resolution. Second, it incorporates an explicit paralogy resolution step that identifies and collapses duplicated linkage groups (arising from WGDs or segmental duplications) into unified pre-duplication representative units prior to ancestral reconstruction, preventing the topology artifacts that confound existing methods. Third, synteny delineation is performed using a Hidden Markov Model (HMM) framework that naturally accommodates biological and technical noise, including gene loss, micro-inversions, and assembly fragmentation, followed by statistical enrichment testing to define high-confidence syntenic blocks. Ancestral reconstruction proceeds via iterative post-order traversal of the species tree, integrating information from all descendant lineages at each node rather than relying on fixed local comparisons. This design reduces sensitivity to lineage-specific genome configurations and allows inference across phylogenies in which no single species provides a stable reference.

**Figure 1.**
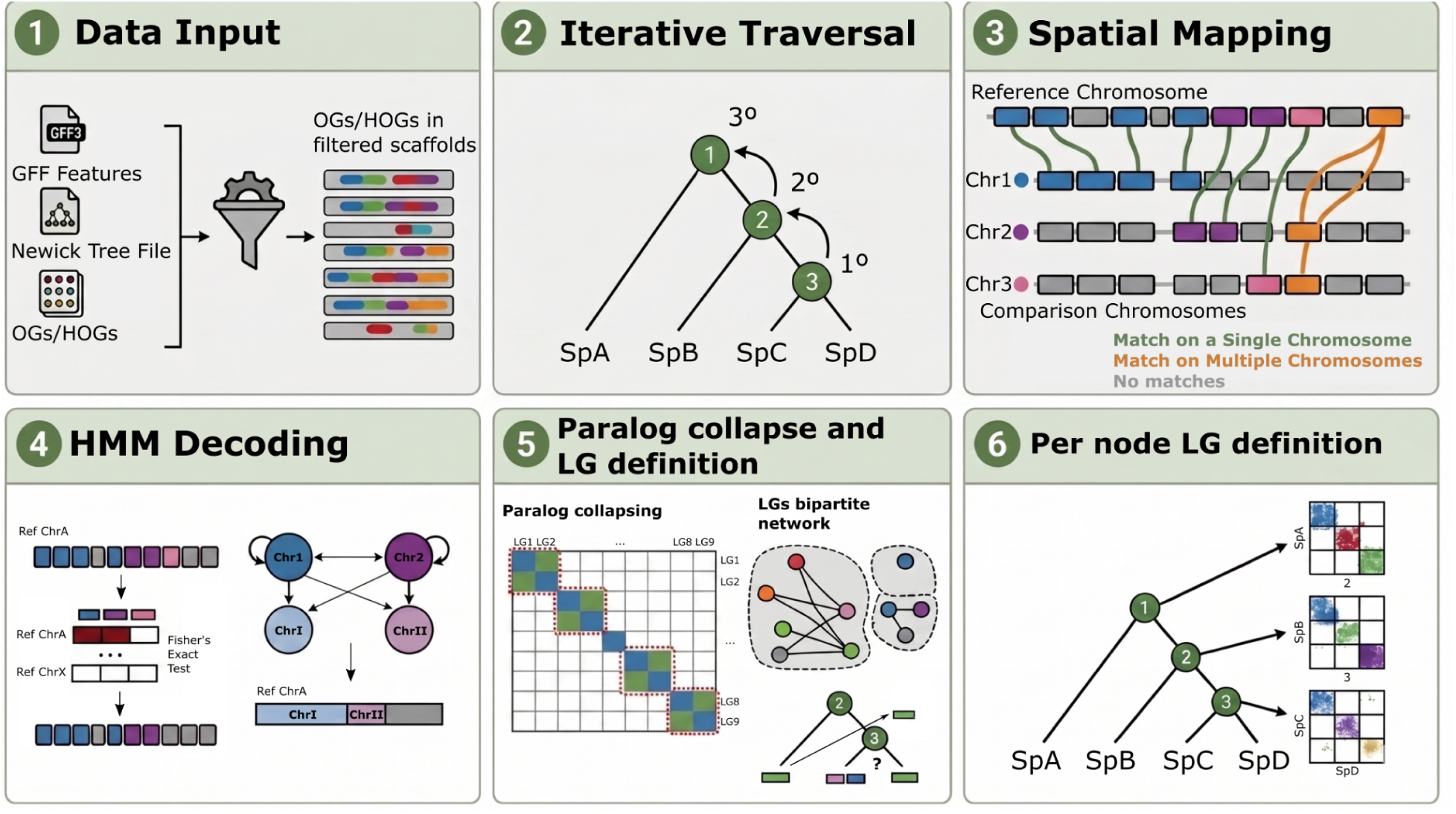
Overview of the LINGUINE pipeline for phylogeny-aware ancestral linkage group inference. LINGUINE takes as input genome assemblies, gene annotations, a rooted species tree, and orthogroup assignments, supporting both OGs and HOGs to maximise marker density. Pairwise synteny is established using a Hidden Markov Model that accommodates gene loss, rearrangements, and assembly fragmentation, followed by statistical enrichment testing to define high-confidence syntenic blocks. An explicit paralogy resolution step collapses duplicated linkage groups arising from whole-genome or segmental duplications prior to reconstruction. Ancestral linkage groups are then inferred iteratively via post-order traversal of the species tree, integrating evidence from all descendant lineages at each node, with outgroup polarisation resolving fusion and fission ambiguities via maximum parsimony.

We validate LINGUINE on nematodes as a gold-standard benchmark, demonstrating performance concordant with existing tools in tractable cases, and apply it to clitellate annelids (a clade with highly rearranged genomes) where LINGUINE recovers biologically coherent ancestral architectures that existing methods fail to resolve. Using GARLIC (Genome reARrangement simuLator for Inferring Chromosomal Landscapes), a purpose-built genome rearrangement simulator, we systematically characterise the limits of synteny signal detectability and define the evolutionary conditions (in terms of rearrangement rate, fusion/fissions events and gene loss) under which ALG reconstruction remains reliable.

## Results

### Benchmarking and sensitivity analysis via stochastic simulations

To define the operational boundaries of LINGUINE and characterise the conditions under which ancestral reconstruction remains reliable, we conducted a systematic sensitivity analysis using GARLIC. Rather than relying solely on empirical benchmarks, where ground truth is always partially unknown (as in the case of the Nigon elements in nematodes and ClitLGs in clitellate annelids, as discussed below), stochastic simulations allow precise control over the evolutionary parameters governing genome evolution, enabling a rigorous assessment of method performance across a wide range of scenarios.

All simulations were based on a three-species topology ((Species A, Species B), Species C), representing the minimal phylogenetic configuration required to perform outgroup-polarised ancestral reconstruction. The simulated ancestral genome comprised 5,000 orthogroups distributed across 15 chromosomes, a scale broadly representative of compact animal genomes. Gene copy number per orthogroup was drawn from a shifted geometric distribution (p = 0.9), ensuring that the majority of orthogroups were single-copy while allowing a realistic proportion of multi-copy families. Gene lengths were set between 500 and 2,000 bp, and a 70% probability was applied to retain paralogous gene copies on the same chromosome, reflecting the empirical tendency of recent duplicates to remain in proximity before dispersal by rearrangement. Full details of the simulation parameters are given in the Methods.

We evaluated two complementary metrics throughout all scenarios: topological accuracy, defined as the correct recovery of ancestral LG number at Node AB (the MRCA of Species A and B), and information retention, defined as the number of orthogroups contributing to reconstructed ALGs. Together, these provide a joint assessment of reconstruction fidelity and the usable genomic signal available for downstream analyses. Four evolutionary scenarios were evaluated, as explained below, each with 100 independent replicates per step.

### Scenario A: Resilience to gene loss

Gene loss is a pervasive feature of genome evolution and a fundamental challenge for synteny-based methods. As lineages diverge and independently lose genes, the pool of shared orthologous anchors available to connect homologous chromosomal regions progressively depletes. To characterise the tolerance of LINGUINE to this process, we simulated increasing levels of independent gene loss in Species A and Species B simultaneously, ranging from 10% to 90% of the orthogroup complement (Fig. 1A), and evaluated the accuracy of ancestral LG inference at Node AB.

LINGUINE maintained perfect topological fidelity, recovering exactly 15 LGs at a median, for gene loss rates between 10% and 60% (Fig. 2B). A sharp transition was observed at 70% loss, where the median inferred LG count increased from 15 to 22, reflecting the failure of the bipartite graph as the number of shared orthogroups per chromosome pair drops below the minimum required for reliable clustering. At this threshold, the ancestral karyotype effectively shatters into numerous gene-poor fragments that can no longer be confidently merged. At 80% and 90% loss, the residual shared signal becomes so critically sparse that even these fragments are eliminated by the pipeline’s noise filter, sweeping the recovered LG count toward zero (Fig. 2B, C). Orthogroup retention follows a concave-down decreasing trajectory across the gradient, with a gradual initial decline that accelerates sharply beyond the 60% threshold (Fig. 2C), providing a measurable and interpretable diagnostic signal that mirrors the topological transition. These results establish 60% independent gene loss as a practical upper bound for reliable reconstruction under this scenario, and identify orthogroup retention as a useful empirical indicator of proximity to this boundary in real datasets.

**Figure 2.**
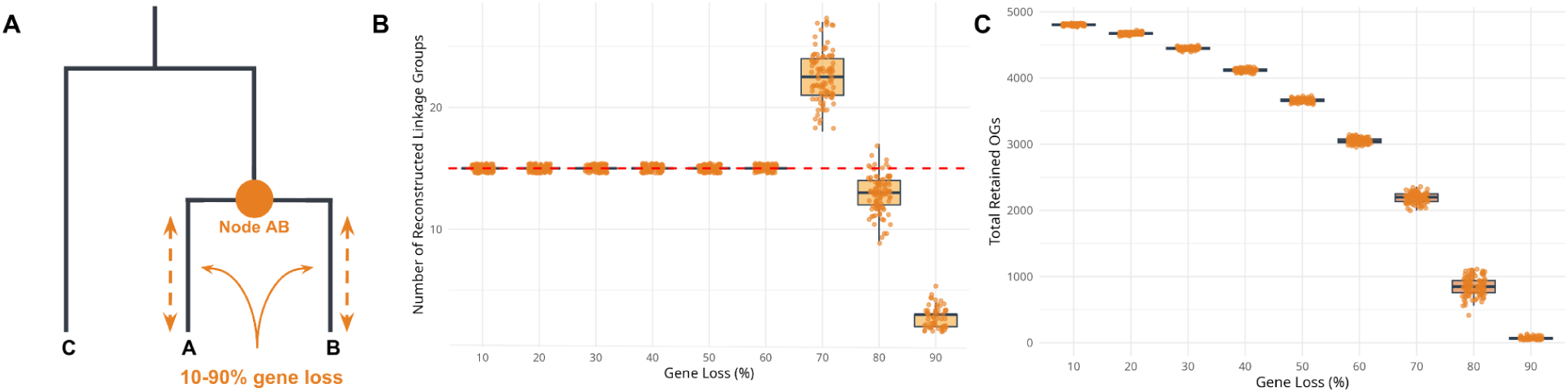
Limits of linkage group inference under increasing levels of independent gene loss. (A) Simulated phylogenetic scenario: independent gene loss of 10% to 90% applied to the branches leading to Species A and Species B. (B) Inferred number of ancestral LGs at Node AB across the gene loss gradient. LINGUINE recovers the correct number of 15 LGs up to approximately 60% loss, followed by overestimation at 70% due to bipartite graph fragmentation, and collapses toward zero at extreme loss levels as the noise filter removes signal-poor fragments. (C) Number of orthogroups retained in reconstructed ALGs under the same conditions. Retention follows a concave-down decreasing trend, reflecting progressive depletion of homologous anchors and providing a practical diagnostic for reconstruction reliability.

### Scenario B: Robustness to post-WGD stochastic gene loss

WGD events present a unique challenge for ancestral reconstruction, as polyploidisation doubles the gene repertoire and distributes homologous blocks across two sets of chromosomes. This difficulty is exacerbated by stochastic gene loss, whereby the independent loss of redundant gene copies in descendant lineages can obscure the shared ancestral signal. We evaluated LINGUINE’s ability to resolve these duplicated states into a unified ancestral configuration. We simulated a WGD event at Node AB, followed by a gradient of independent random gene loss in both daughter lineages ranging from 10% to 90% (Fig. 3A).

**Figure 3.**
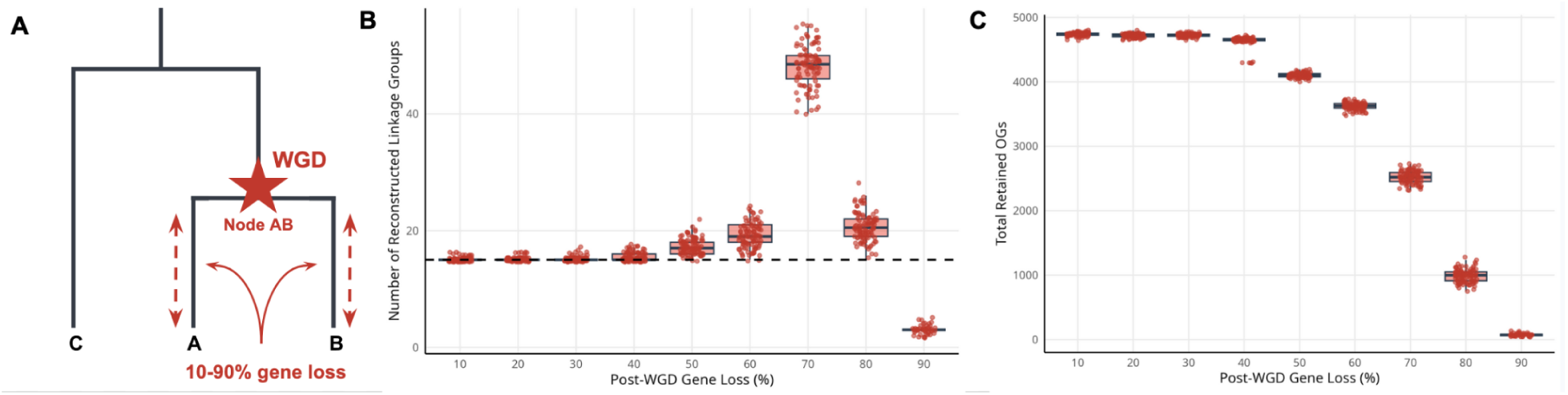
Limits of linkage group inference under post-WGD stochastic gene loss. (A) Species tree describing the simulated events (i.e. a WGD event happening in the MRCA of species A and B). (B) Inferred number of ancestral linkage groups following a simulated WGD event at Node AB and increasing levels of independent gene loss in descendant lineages (10–90%). LINGUINE accurately collapses duplicated chromosomal complements up to ∼40% gene loss, with a sharp transition to overestimation of linkage group number beyond ∼70%. (C) Number of orthogroups retained in reconstructed ALGs under the same conditions. Orthogroup retention follows a concave-down decreasing trend, starting with a more gradual decline followed by a rapid loss at 70% gene loss reflecting progressive depletion of homologous anchors required for paralogy resolution.

The resulting fragmentation curve was virtually identical to that from the gene loss scenario (Scenario A; Fig. 3B and 3C), demonstrating that LINGUINE successfully collapses duplicated genomic content into the correct ancestral units despite the severe removal of orthologous anchors. This shows that LINGUINE can profit from paralogs and ohnologs, which are routinely discarded by other methodologies, to reconstruct topology.

### Scenario C: Robustness to outgroup signal erosion and structural remodelling via fusion and fission events

Resolving ancestral karyotypes frequently requires polarising structural ambiguities, particularly when distinguishing between lineage-specific chromosome fusions and fissions. These two processes produce complementary signals that are indistinguishable without external context: a derived fission in one lineage and a derived fusion in another leave an identical imprint on the pair-wise comparison between ingroups. To evaluate LINGUINE’s capacity to resolve this class of ambiguity, and to establish how much outgroup information is actually required, we designed two complementary simulations based on the same underlying phylogenetic scenario.

We first modelled a karyotypic divergence in which Species B underwent a lineage-specific fission event (yielding 16 chromosomes) relative to Species A, which retained the ancestral state of 15 chromosomes (Fig. 4A). Without external context, both an ancestral state of 15 chromosomes followed by fission in B, and an ancestral state of 16 chromosomes followed by fusion in A, are equally parsimonious. LINGUINE resolves this symmetry by querying syntenic evidence from the outgroup Species C, using outgroup-based polarisation to determine the ancestral configuration via maximum parsimony.

**Figure 4.**
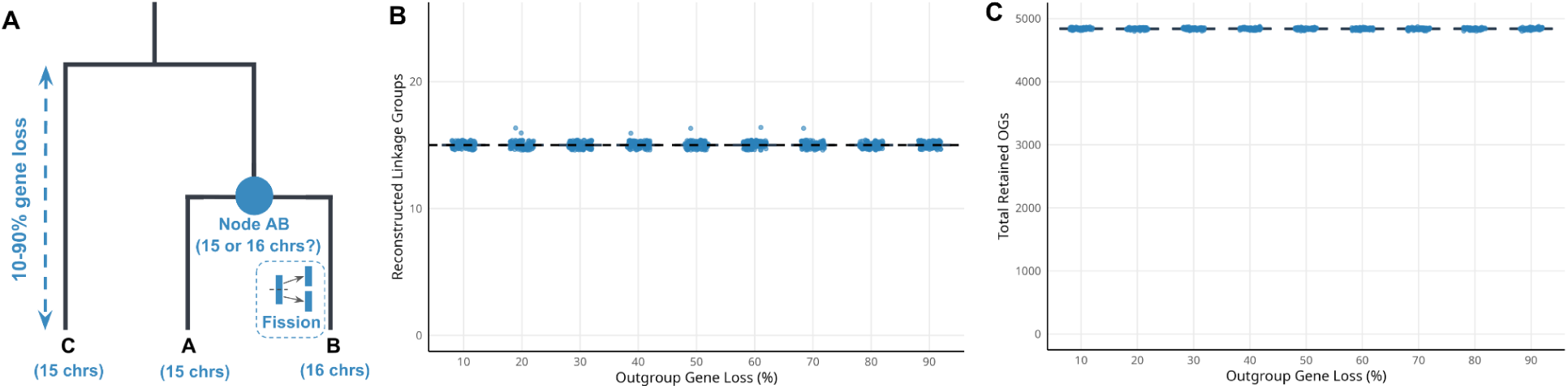
Robustness of outgroup-based polarisation under progressive outgroup signal erosion. (A) Species tree describing the simulated scenario. Species B has 16 chromosomes due to a lineage-specific fission, while Species A retains the ancestral complement of 15. Two equally parsimonious ancestral states exist without outgroup evidence. (B) Reconstruction accuracy at Node AB under increasing gene loss in the outgroup (Species C; 10–90%). LINGUINE correctly resolves the 15 LG ancestral state in 99.2% of replicates across the full gradient, demonstrating that polarisation is robust to severe outgroup signal erosion. (C) Orthogroup retention in reconstructed ALGs across the same gradient. Retention remains highly stable, indicating that only a fractional outgroup signal is required for reliable ancestral state inference.

To establish the minimum outgroup signal required for reliable polarisation, we simulated a gradient of random gene loss in Species C alone (10% to 90%), leaving the ingroup genomes intact. LINGUINE maintained near-perfect reconstruction accuracy across the entire gradient, achieving a 99.2% accuracy rate (893/900 replicates) and consistently recovering the correct 15 LG ancestral state (Fig. 4B). The rare deviations from this expectation (fewer than 1% of replicates) were entirely stochastic and showed no correlation with the severity of outgroup gene loss. The number of orthogroups retained in reconstructed ALGs remained highly stable across all loss levels (Fig. 4C), confirming that even a fractional outgroup signal is sufficient to break topological ambiguity reliably. This robustness is particularly important for empirical datasets, where outgroup lineages may be phylogenetically distant or evolving at elevated rates.

We next asked how LINGUINE performs when the ingroup itself has experienced extensive chromosomal restructuring, even in the absence of net changes in chromosome number. Sequential fusion and fission events present a distinct and more difficult challenge, as they progressively erode the syntenic identity of extant chromosomes relative to their ancestral progenitors while preserving overall chromosome count, effectively concealing the true magnitude of structural remodelling from karyotype-level comparisons. To model this, we simulated 5 to 45 matched pairs of fission and fusion events in Species A (Fig. 5A). This balanced design extensively reshuffled gene composition and internal chromosomal macrostructure without altering the number of chromosomes, creating a scenario in which the genomic architecture has been substantially scrambled but no superficial karyotypic signal of rearrangement remains.

**Figure 5.**
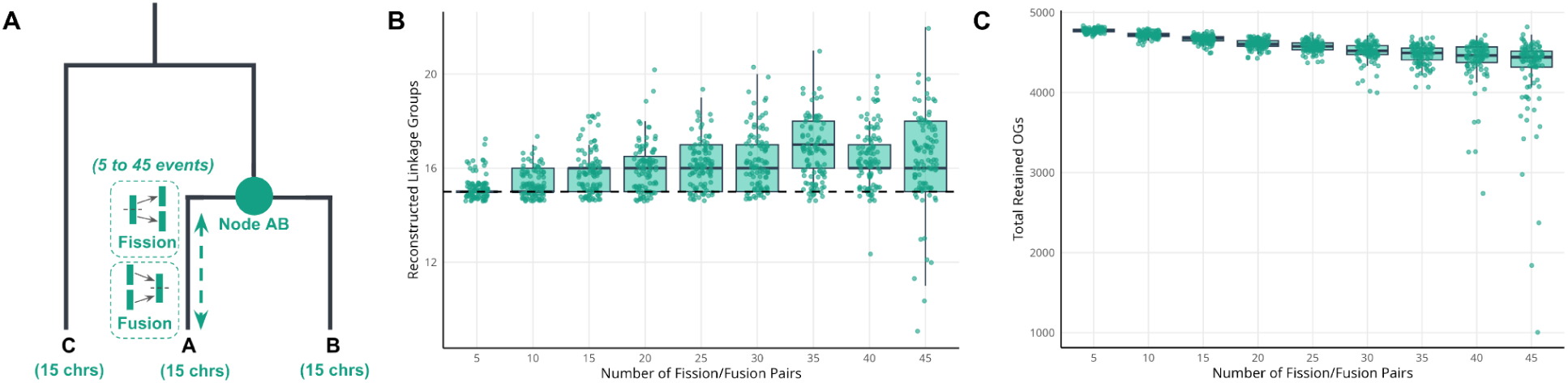
Resilience of LINGUINE to structural remodeling via fusion and fission events. (A) Species tree describing the simulated events. (B) Inferred number of ancestral linkage groups under increasing numbers of matched fusion and fission events (5–45 pairs) in Species A. Despite extensive reshuffling of gene composition, LINGUINE predicts a number of LGs similar to the correct ancestral chromosome number (15 LGs), although with an increasing variance. (C) Number of orthogroups retained in reconstructed ALGs across the same simulations. Orthogroup retention remains stable, indicating that rearrangements that redistribute genes in large blocks preserve sufficient information for accurate reconstruction.

LINGUINE demonstrated exceptional resilience under these conditions. The mean recovered LG count shifted only modestly across the full rearrangement gradient, from 15.2 at 5 pairs to a maximum of 16.7 at 35 pairs (Fig. 5B). At the most extreme threshold (45 pairs), variance increased substantially, spanning 9 to 22 inferred LGs, reflecting a conservative tendency to isolate highly scrambled sub-fragments rather than erroneously merge distinct ancestral chromosomes. Orthogroup retention remained largely stable across the gradient, declining gradually from approximately 4,774 to 4,271 OGs (Fig. 5C), indicating that large-block rearrangements preserve sufficient inter-species syntenic linkage for the bipartite graph to function correctly. Taken together, these results demonstrate that LINGUINE’s outgroup polarisation and graph clustering logic are jointly robust to both the loss of outgroup information and to extensive ingroup structural remodelling, two of the most common sources of uncertainty in real-world ancestral reconstruction problems.

### Scenario D: Resilience to inter-chromosomal translocations

Inter-chromosomal translocations are among the most pervasive drivers of chromosomal evolution across animal lineages, and represent a particularly challenging signal to interpret in the context of ancestral reconstruction. Unlike fusions and fissions, which alter chromosome number and leave a detectable karyotypic footprint, translocations redistribute gene content across chromosomes while preserving overall chromosome count, progressively dissolving the boundaries between ancestral linkage groups without any superficial signal of rearrangement. As translocation events accumulate, syntenic blocks are broken into smaller and more dispersed fragments, reducing the density of contiguous orthologous anchors available to connect extant chromosomal regions to their ancestral progenitors.

To evaluate LINGUINE’s tolerance to this class of rearrangement, we simulated a gradient of 50 to 450 non-reciprocal inter-chromosomal translocation events in Species A, applied iteratively to mimic the cumulative nature of chromosomal evolution over time. Each event involved the relocation of a contiguous gene block of 5 to 40 genes (Fig. 6A), producing complex nested rearrangements with increasing depth as events accumulated. Species B and Species C were left unmodified throughout, so that any degradation in reconstruction accuracy could be attributed directly to the rearrangement burden in Species A.

**Figure 6.**
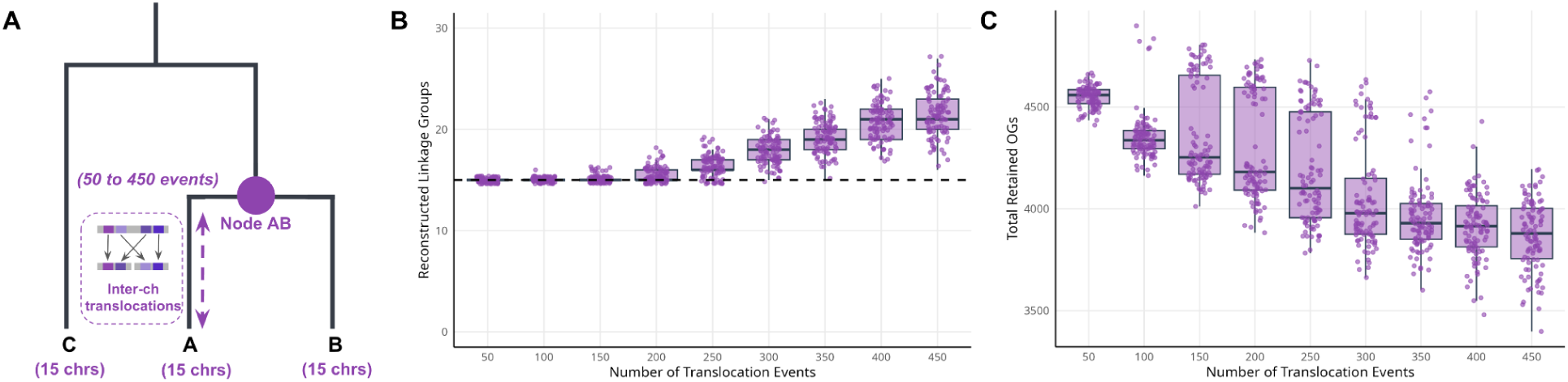
Robustness of LINGUINE to inter-chromosomal translocation accumulation. (A) Simulated phylogenetic scenario: 50 to 450 non-reciprocal inter-chromosomal translocation events applied iteratively to Species A, each relocating a contiguous block of 5 to 40 genes. (B) Inferred number of ancestral LGs at NodeAB across the translocation gradient. LINGUINE maintains accurate reconstruction up to approximately 200 cumulative events, beyond which the inferred LG count increases progressively, reflecting conservative fragmentation of sub-threshold syntenic blocks rather than erroneous merging of distinct chromosomes. (C) Number of orthogroups retained in reconstructed ALGs across the same gradient. Retention declines progressively with increasing translocation load, reflecting the erosion of contiguous syntenic signal and the reduction of recoverable ancestral segments.

LINGUINE maintained perfect topological fidelity, recovering a mean of exactly 15 inferred LGs, for up to 200 cumulative translocation events (Fig. 6B). This represents a substantial level of chromosomal disruption: at 200 events, with blocks of up to 40 genes relocated per event, a significant fraction of ancestral chromosomal gene content has been redistributed across non-homologous chromosomes. A transition to over-fragmentation was observed at 250 translocations, where the median inferred LG count rose to 16, increasing progressively to a median of 21 at the highest rearrangement level tested. This upward drift reflects the progressive breakdown of contiguous syntenic blocks below the minimum size required for reliable clustering, rather than errors in the fundamental logic of the reconstruction. The pipeline does not erroneously merge distinct chromosomes but instead conservatively isolates sub-fragments whose ancestral identity can no longer be confidently assigned.

Orthogroup retention declined in parallel with topological accuracy, decreasing from a median of 4,559 OGs at 50 translocation events to 3,880 at 450 events (Fig. 6C). This progressive decline reflects the dual consequence of translocation accumulation: not only does the reconstructed architecture become increasingly fragmented, but the informative gene repertoire itself contracts as small ancestral segments become too dispersed to be recovered as coherent blocks. The two metrics therefore provide complementary and consistent diagnostics of reconstruction reliability across this rearrangement gradient.

Taken together, these results establish approximately 150 to 200 cumulative non-reciprocal translocations as the practical upper bound for reliable reconstruction in genomes of this size and configuration. Beyond this threshold, degradation is progressive and measurable rather than abrupt, providing a useful empirical signal for identifying when reconstruction results should be interpreted with caution in real datasets. Importantly, the capacity of LINGUINE to buffer up to 200 nested translocations while maintaining accurate ancestral LG inference substantially exceeds what would be achievable using sparse single-copy marker sets, underscoring the practical advantage of orthogroup-based signal in highly rearranged genomes.

### Reconstruction of the ancestral nematode karyotype

Nematodes offer a well-suited benchmark for ALG reconstruction methods. Their genomes evolve rapidly, accumulating extensive gene order rearrangements, yet their ancestral chromosomal organisation is precisely characterised by seven conserved elements, the Nigon elements A, B, C, D, E, N, and X, whose content and boundaries are documented and carefully curated across multiple independent lineages (Tandonnet et al. 2019; Gonzalez de la Rosa et al. 2021). This provides a well-supported reference set against which both topological accuracy and gene assignment specificity can be directly evaluated.

We applied LINGUINE to a dataset of 13 nematode genomes spanning Rhabditina and Tylenchina (Fig. 7A) and compared its performance to odp and Syngraph, which were run in the same dataset (see Methods). A fundamental distinction among these methods lies in their capacity to infer ancestral states at internal nodes rather than simply describing shared linkage among extant species. Odp operates as a purely pairwise framework and, while useful for identifying conserved chromosomal correspondences, cannot reconstruct independent ancestral configurations at internal nodes. Its output at node N1, the MRCA of Rhabditina and Tylenchina, was consequently a highly fragmented set of over 100 small linkage groups with little biological interpretability, reflecting the inability of pairwise comparison to integrate syntenic signal distributed across multiple rearranged genomes (Fig. 7C). Syngraph, which does implement node-by-node ancestral reconstruction, recovered a broadly correct number of linkage groups but failed to fully separate the N and X Nigon elements, instead depicting them as a partially fused unit. LINGUINE correctly recovered all seven expected linkage groups at node N1, cleanly resolving N and X as distinct ancestral units (Fig. 7C).

**Figure 7.**
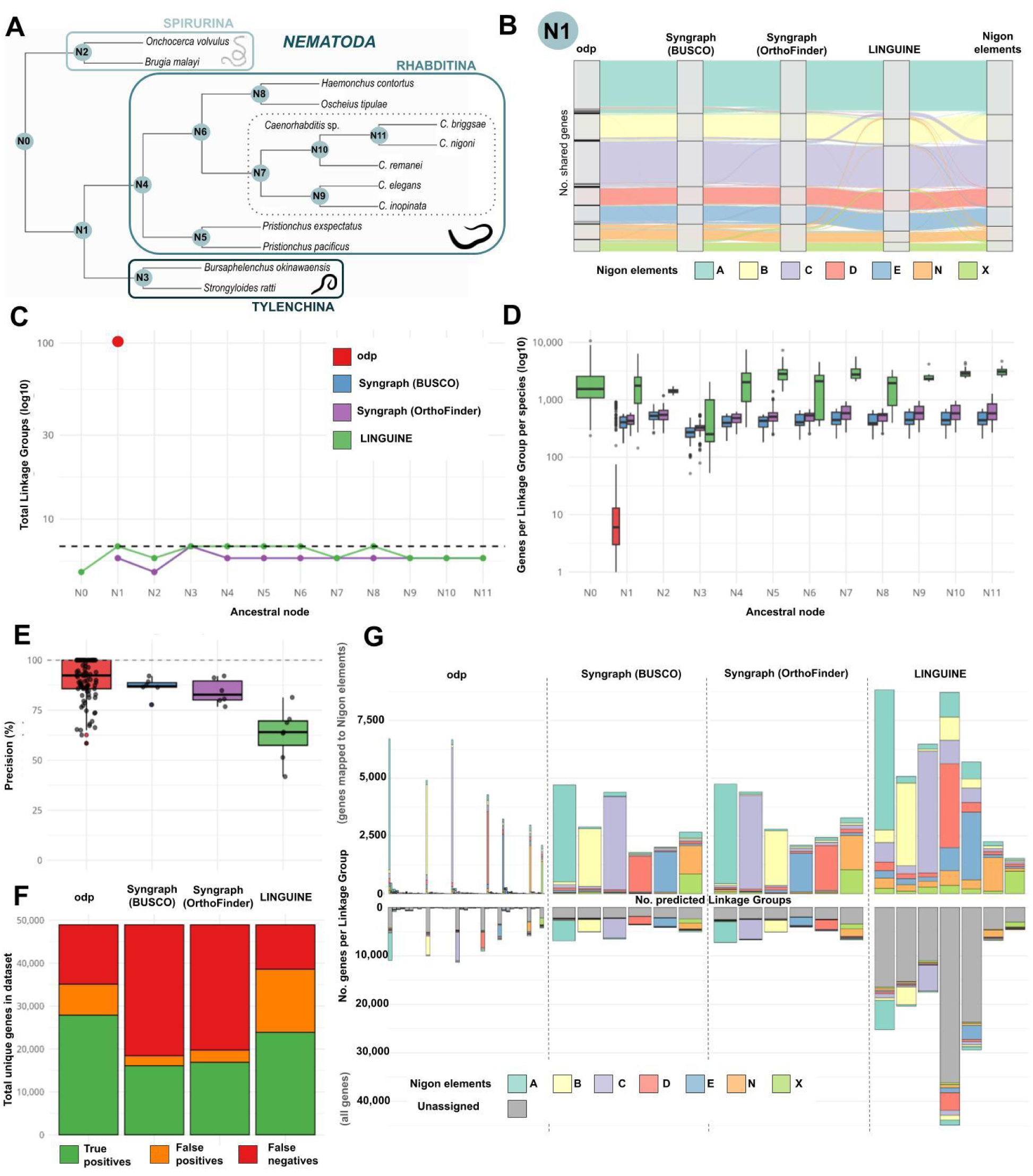
Reconstruction of the ancestral nematode karyotype. (A) Species phylogeny of the 13 nematode genomes used for ancestral reconstruction. Node N1 indicates the most recent common ancestor of Rhabditina and Tylenchina, the primary node used for benchmarking. (B) Alluvial plot of gene assignments across methods and Nigon elements. Each thread represents one gene, coloured by its Nigon element classification. Only genes assigned by all three methods and with a known Nigon element classification are shown (n = 240). Shared misassignments across methods reflect intrinsic signal ambiguity rather than method-specific errors. (C) Number of inferred ancestral linkage groups across all reconstructed nodes for each method. The dashed line indicates the expected seven Nigon elements. LINGUINE recovers seven linkage groups at node N1 and correctly separates the N and X elements as distinct units. Syngraph recovers a comparable count but shows residual fusion of N and X. Odp produces a highly fragmented output of over 100 small groups with limited biological interpretability. (D) Number of genes assigned to reconstructed linkage groups per method. LINGUINE incorporates a substantially larger fraction of the gene repertoire by using hierarchical orthogroups, whereas Syngraph is largely restricted to conserved single-copy markers. (E) Per-element precision (proportion of genes in each reconstructed LG that map to the correct Nigon element) across methods. Smaller linkage groups tend to achieve higher precision scores. LINGUINE shows somewhat lower median precision than Syngraph, reflecting the broader and noisier orthogroup signal. (F) Classification of all genes assigned to a Nigon element: true positives (correct Nigon assignment), false positives (incorrect Nigon assignment), and false negatives (genes not assigned to any LG). LINGUINE recovers a substantially greater number of true positives in absolute terms. (G) Genome-wide correspondence between reconstructed linkage groups and Nigon elements across methods. Upper panel: correspondence for all genes assigned to an LG. Lower panel: same, extended to include genes not assigned to any LG. LINGUINE achieves broader genome coverage at a modest cost to per-gene specificity.

A second and equally important distinction concerns marker coverage. By leveraging HOGs rather than conserved single-copy BUSCO markers, LINGUINE incorporates a substantially larger fraction of the total gene repertoire into the reconstruction (Fig. 7D). This broader signal enables syntenic inference across genomic regions that are entirely excluded from marker-restricted analyses, and allows LINGUINE to reconstruct coherent ancestral linkage groups at deeper nodes in the phylogeny, including the root of the 13-species tree, where the sparsity of conserved single-copy markers becomes a limiting factor for competing methods.

This increased coverage comes with a documented trade-off in assignment precision. When evaluated against the canonical Nigon element assignments, LINGUINE shows slightly lower per-element precision than Syngraph (Fig. 7E), reflecting the inclusion of lineage-specific gene family expansions, multi-copy orthogroups, and rapidly evolving genes that carry inherently ambiguous chromosomal signals. It is important to note, however, that this comparison is made across a very large disparity in the number of genes used: LINGUINE operates on a gene set that is substantially larger than the conserved single-copy marker set available to Syngraph (Fig. 7D). Despite this, LINGUINE correctly classifies a greater absolute number of genes as true positives across all seven Nigon elements (Fig. 7F), demonstrating that broader marker inclusion translates into a net gain in correctly assigned genes, not merely a redistribution of error. Odp, while achieving a high number of true positives in absolute terms, does so by distributing genes across a very large number of small and biologically uninterpretable linkage groups (Fig. 7C, 7F), which severely limits the utility of its output for downstream ancestral reconstruction. Notably, a consistent subset of misassignments was shared across all three methods (Fig. 7B), pointing to intrinsic ambiguity in the syntenic signal itself rather than method-specific artifacts, and setting a biological ceiling on the precision achievable by any approach. Together, these results establish a clear and interpretable performance profile: LINGUINE sacrifices a modest degree of per-assignment specificity to achieve substantially broader and more complete reconstruction of the ancestral genome (Fig. 7G).

### Application to clitellate annelids reveals signal recovery in highly rearranged genomes

Having established LINGUINE’s performance profile in a system with a known ancestral karyotype, we next applied it to clitellate annelids, a clade where the evolutionary forces that challenge existing methods operate at their most extreme. Clitellates exhibit extensive interchromosomal rearrangements, pronounced lineage-specific variation in chromosome number, and probable polyploidisation events in multiple lineages, making their ancestral chromosomal architecture challenging to marker-restricted approaches (Lewin et al. 2024; Vargas-Chávez et al. 2025). The availability of several chromosome-scale assemblies and an established set of manually curated clitellate linkage group reference elements (named ClitLGs) originally defined in Vargas-Chávez et al. (2025) (Vargas-Chávez et al. 2025) using pairwise odp comparisons followed by manual curation based on Fisher’s exact tests, makes this an informative empirical system in which to assess real-world reconstruction performance beyond a controlled benchmark.

LINGUINE recovered coherent and stable ancestral linkage groups across all internal nodes of the clitellate phylogeny, including the root (Fig. 8A, C). Strikingly, the ALGs inferred by LINGUINE are highly congruent with the ClitLG elements defined in (Vargas-Chávez et al. 2025), despite the two approaches being methodologically entirely independent: while that study relied on pairwise synteny comparisons with manual expert curation, LINGUINE reconstructs ancestral states automatically through phylogeny-wide graph clustering with no manual intervention. This congruence provides strong cross-validation for both the ClitLG elements themselves and for LINGUINE’s automated reconstruction logic, and demonstrates that the ancestral chromosomal signal in clitellates is robust enough to be recovered by fundamentally different analytical strategies.

**Figure 8.**
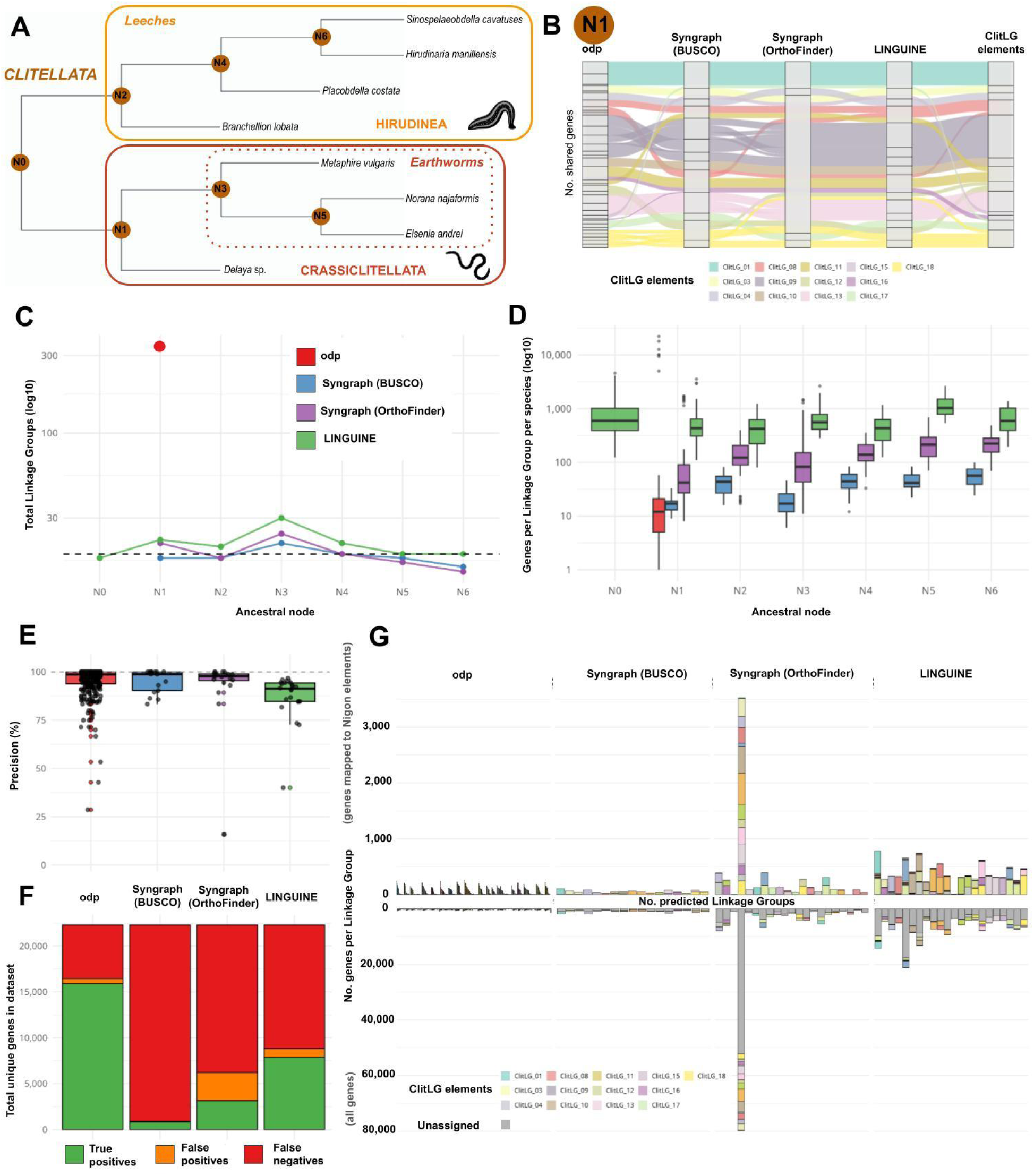
Application of LINGUINE to clitellate annelids reveals recovery of ancestral chromosomal structure in a highly rearranged clade. (A) Species phylogeny used for ancestral reconstruction. (B) Inferred ancestral linkage groups across all internal nodes of the clitellate phylogeny. LINGUINE recovers coherent and stable linkage groups throughout the tree despite extensive chromosomal rearrangement in extant lineages. (C) Comparison of gene coverage and linkage group structure across methods. Odp produces highly fragmented outputs; Syngraph recovers more coherent groups but with limited gene coverage; LINGUINE integrates a substantially larger fraction of the gene repertoire, enabling signal recovery where other methods fail. (D) Per-element assignment precision against established ClitLG reference elements across methods. (E) Classification of genes assigned to a ClitLG element as true positives, false positives, or false negatives. LINGUINE recovers the greatest number of true positives in absolute terms. (F) Correspondence between reconstructed LGs and ClitLG elements for genes with confirmed ClitLG membership. (G) Same correspondence extended to all classified genes. (H) Alluvial plot showing assignment consistency for genes grouped into an LG by all methods and classified into a ClitLG element; the limited gene count reflects the restricted marker set enforced by BUSCO-based approaches.

In contrast, odp produced highly fragmented outputs consistent with the well-characterised failure mode of pairwise approaches in extensively rearranged systems. Syngraph recovered a more coherent set of linkage groups but was substantially constrained by its sparse marker coverage, limiting its ability to detect syntenic signal across the broad fraction of the genome where conserved single-copy orthologs are absent (Fig. 8D). The relative performance advantage of LINGUINE over both methods is considerably more pronounced here than in nematodes, and for a principled reason: in highly rearranged genomes, the primary obstacle to reconstruction is not assignment precision but the availability of sufficient shared markers to detect macrosyntenic signal in the first place. By incorporating a broader orthogroup-based signal and integrating evidence across the full phylogeny, LINGUINE recovers chromosomal structure that is effectively invisible to approaches constrained by sparse marker sets.

Assessment of the reconstructed LGs against the established ClitLG reference elements confirmed the coverage-precision trade-off observed in nematodes. LINGUINE shows slightly lower per-element assignment precision than Syngraph (Fig. 8E), as expected from the inclusion of multi-copy orthogroups and lineage-specific gene families. However, the total number of genes correctly classified as true positives is substantially higher for LINGUINE than for either comparator (Fig. 8F), reflecting the practical value of its broader genomic coverage in a clade where marker density is the binding constraint. The correspondence between

LINGUINE’s reconstructed ALGs and the ClitLG elements is well maintained both when the comparison is restricted to genes with confirmed ClitLG membership and when extended to the full classified gene set (Fig. 8G), indicating that the broader orthogroup signal does not introduce systematic misassignment but rather expands the informative gene repertoire while preserving overall structural fidelity. The alluvial plot in Fig. 8B illustrates assignment consistency for the subset of genes recovered by all three methods simultaneously. The limited size of this shared set reflects the fundamental constraint imposed by BUSCO-based approaches, which exclude the majority of the gene repertoire from consideration and thereby restrict the scope of any cross-method comparison.

## Discussion

Reconstructing the ancestral chromosomal complement of extinct lineages is a challenging task in comparative genomics, and its difficulty scales with the very features that make it most biologically interesting. In clades where genome evolution has been fast, high rates of chromosomal rearrangement erode syntenic continuity, lineage-specific gene loss depletes the shared orthologous markers that underpin reconstruction, and WGD events complicate the interpretation of paralogy. Existing methods address this problem with tools that were largely designed for tractable, slowly evolving systems: sparse single-copy markers, fixed local comparisons, and no mechanism to resolve duplicated chromosomal content (Mackintosh et al. 2023; Schultz et al. 2023; El Hinani & Copley 2023). LINGUINE was built to operate where these approaches break down, combining orthogroup-based marker expansion, explicit paralogy resolution, and phylogeny-aware reconstruction that integrates evidence across the full species tree.

A central contribution of this work is the systematic characterisation of the conditions under which ancestral reconstruction remains reliable. We used GARLIC to sweep key evolutionary parameters across wide ranges under fully controlled conditions. The results reveal a consistent pattern: LINGUINE maintains high topological accuracy up to approximately 60% independent gene loss, up to 200 cumulative inter-chromosomal translocation events, and across the full range of matched fusion and fission rearrangements tested, before entering a regime of progressive over-fragmentation. Crucially, these transitions are gradual and are accompanied by measurable declines in orthogroup retention, providing practical diagnostics that can be evaluated in empirical datasets to assess whether a given reconstruction is operating within the reliable range. The near-complete robustness to outgroup signal erosion is a particularly important result: even outgroups retaining only a small fraction of shared markers were sufficient to correctly polarise ambiguous ancestral states in 99.2% of replicates, a finding with direct practical implications given that real outgroup lineages are frequently distant, fragmented, or rapidly evolving.

Benchmarking on nematodes places LINGUINE’s performance in the context of a well-characterised biological system. The correct recovery of all seven Nigon elements, including clean separation of N and X, and the reconstruction of coherent ancestral states across all nodes of the 13-species phylogeny confirm that LINGUINE performs at or above the level of existing tools in tractable cases. The principal trade-off is one of precision versus coverage. By incorporating HOGs rather than single-copy markers, LINGUINE assigns genes to ancestral LGs across a substantially broader fraction of the genome, but the inclusion of multi-copy orthogroups, lineage-specific gene family expansions, and rapidly evolving genes introduces assignment noise that reduces per-element precision relative to marker-restricted approaches. Critically, this precision cost is more than offset by the gain in absolute true positive assignments, as LINGUINE correctly classifies more genes in total than either comparator, even accounting for the much larger gene set it operates on. The observation that a consistent subset of misassignments is shared across all three methods suggests that these discrepancies do not reflect method-specific errors but rather point to limitations in the Nigon elements themselves as a reference framework (Tandonnet et al. 2019). As chromosome-scale assemblies continue to accumulate across nematode diversity, the boundaries and gene membership of these elements will likely need to be revisited and refined, and the shared misassignments reported here may provide a useful starting point for that effort.

The application to clitellate annelids reveals the practical value of LINGUINE’s design in a system where the limitations of existing methods become acute. In this clade, extensive chromosomal rearrangement and probable polyploidisation events in multiple lineages reduce the macrosyntenic signal available to any marker-restricted approach below the threshold required for coherent reconstruction (Lewin et al. 2024; Vargas-Chávez et al. 2025). Under these conditions, odp yields highly fragmented outputs and Syngraph recovers a more coherent structure but remains constrained by sparse marker coverage. LINGUINE recovers stable and interpretable ancestral linkage groups across all internal nodes of the phylogeny, including the root, and these ALGs are highly congruent with the ClitLG elements originally defined in (Vargas-Chávez et al. 2025) through an entirely independent methodology. This cross-method agreement is a strong indication that the recovered ancestral architecture reflects genuine biological signal rather than analytical artefact. The relative performance advantage of LINGUINE over comparator methods is more pronounced in clitellates than in nematodes: as the availability of shared orthologous markers becomes the primary limiting factor for signal detection, the benefit of orthogroup-based inference over single-copy approaches grows proportionally, suggesting that LINGUINE will provide the greatest gains precisely in the systems where it is most needed.

A key feature underlying this behaviour is the phylogeny-aware reconstruction strategy. By performing iterative reconstruction through post-order traversal of the species tree and progressively integrating genomic information from all descendant lineages at each node, LINGUINE avoids the dependence on any single reference genome or fixed three-species window that constrains competing methods. This design is particularly important in datasets where no individual species provides a stable or representative chromosomal configuration, a situation that is the rule rather than the exception in rapidly evolving clades.

Several limitations should be acknowledged. Like all synteny-based approaches, LINGUINE depends on the quality of genome assemblies, gene annotations, and orthology inference, and errors in any of these inputs propagate into syntenic block detection and ancestral reconstruction. The method cannot recover signal that has been fully eroded by extreme rearrangement or catastrophic gene loss, as the simulations clearly define. That said, the simulations also demonstrate that LINGUINE tolerates a substantial degree of genomic disruption before reconstruction accuracy degrades: the correct ancestral karyotype is recovered across the full range of fusion and fission events tested, and topological fidelity is maintained up to 150 to 200 cumulative inter-chromosomal translocations, a level of rearrangement that substantially exceeds what is achievable by marker-restricted approaches. These thresholds are relevant for real datasets, as they establish that LINGUINE operates reliably across a wide range of genome evolution rates and rearrangement histories encountered in practice. Inferred linkage groups should therefore always be evaluated alongside orthogroup retention statistics, block continuity, and cross-lineage consistency. The paralogy resolution framework provides strong evidence for large-scale duplication events, but definitive conclusions about WGD timing and mechanism should integrate complementary analyses, including gene tree reconciliation and synonymous divergence profiling.

As chromosome-scale genome assemblies continue blooming across the animal tree of life, the need for tools capable of ancestral genome reconstruction in complex and poorly conserved clades will only grow. LINGUINE fills an important gap in the current methodological toolkit. Together with GARLIC, it provides both the reconstruction framework and the simulation infrastructure needed to interpret empirical results in their proper evolutionary context, and to identify in advance the conditions under which those results can be trusted.

## Materials and Methods

### Input requirements and genomic preprocessing

The pipeline requires reference genome assemblies (FASTA format) and corresponding gene annotations (GFF3 format) for all extant species included in the analysis. Prior to synteny inference, gene coordinates are extracted from GFF3 files and each gene is assigned to its chromosomal location based on the scaffold or chromosome identifier in the assembly. To reduce noise from fragmented assemblies, users can exclude unanchored scaffolds and contigs below a configurable length threshold from all downstream analyses; only sequences above this threshold are retained for synteny delineation and ancestral reconstruction. Orthology inference must be provided as input. While the pipeline is natively optimised for OrthoFinder outputs and supports both OGs and HOGs, its modular architecture accepts orthology assignments from any source in a standard tabular format, provided that gene identifiers match those in the corresponding GFF3 annotation. A rooted species tree in Newick format is required to guide the post-order traversal (from tips to root) used for iterative ancestral reconstruction. To ensure computational stability and prevent memory leakage during large multi-species runs, the orchestrator script dynamically isolates the execution environment at each node of the phylogeny, spawning and terminating R subprocesses on a per-node basis.

### HMM-based synteny delineation

Pairwise synteny between lineages is established using a Hidden Markov Model (HMM) implemented via the HMM R package (https://cran.r-project.org/web/packages/HMM/index.html). For each pairwise comparison, genes of the reference species are ordered by their genomic coordinates along each chromosome or scaffold. The hidden states of the model represent the ancestral chromosomes or LGs, and the observation sequence consists of the chromosomal assignments of orthologous genes from the target species, mapped onto the coordinate framework of the reference.

Transition probabilities are configured to strongly penalise state switching between target chromosomes, allowing the model to bridge localised assembly errors, gene loss, or micro-inversions without fragmenting syntenic tracts (default self-transition probability P(S_i to S_i) = 0.995). Emission probabilities are parameterised to accommodate several sources of biological and technical noise: missing orthologs arising from gene loss or assembly gaps are modelled as uninformative emissions; ambiguous multi-mapping due to paralogy is handled by distributing emission probability across candidate target chromosomes; and low-confidence chromosomal assignments are downweighted. All HMM parameters are exposed to the user through a central configuration file, enabling synteny stringency to be tuned according to the evolutionary distances and assembly qualities of the species under comparison.

Following Viterbi decoding of the optimal state path, a sliding window approach smooths the inferred assignment along each reference chromosome, resolving isolated state switches that are more likely to reflect noise than true rearrangements. Syntenic blocks are then statistically validated using a one-sided Fisher’s Exact Test applied to a contingency table of observed versus expected ortholog co-occurrence (default significance threshold p < 0.01, minimum odds ratio > 2.0). Contiguous runs of statistically significant assignments are collapsed into Strict Blocks, representing high-confidence syntenic units. Strict Blocks mapping to the same target chromosome are subsequently merged into Broad Regions, rescuing intervening genes that lack clear orthology but reside within otherwise syntenic tracts. For comparisons between extant tip species, reciprocal mapping is performed automatically, running the HMM in both directions and merging the resulting syntenic evidence to capture signal from both genomic perspectives.

### Paralogy resolution and background noise filtering

Prior to ancestral reconstruction, the pipeline performs an explicit paralogy resolution step to prevent duplicated linkage groups arising from whole-genome or segmental duplications from introducing artifacts into the bipartite graph topology. An intra-specific all-by-all syntenic comparison is performed across the broad syntenic regions of each species, constructing a chromosome-by-chromosome overlap matrix based on shared orthogroup content. LG pairs within the same species that share a statistically significant orthogroup overlap (Fisher’s Exact Test, Benjamini-Hochberg adjusted p < 1×10^-20^) are flagged as candidate duplicated units and collapsed into unified pre-duplication representative blocks using graph-based clustering implemented via the igraph R package (https://cran.r-project.org/web/packages/igraph/index.html). The resulting merged blocks replace the original duplicated LGs as inputs to the reconstruction step, preventing post-duplication chromosomal copies from being propagated as independent ancestral units.

Additionally, to prevent highly conserved or ubiquitous gene families from generating spurious cross-chromosomal linkages that inflate syntenic connectivity, orthogroups present above a user-defined abundance threshold across LGs are dynamically filtered from the analysis prior to graph construction. This step removes highly dispersed gene families and other broadly distributed orthogroups whose chromosomal distribution carries no informative syntenic signal.

### Ancestral genome reconstruction via graph clustering

Ancestral linkage groups are reconstructed iteratively at each internal node of the species tree, proceeding in post-order from the tips to the root. At each node, a bipartite graph is constructed in which vertices represent the LGs of the two daughter lineages (either extant tip species or previously reconstructed internal nodes), and weighted edges are drawn between LG pairs based on the number of shared orthogroups, after filtering. Edge significance is assessed using a Fisher’s Exact Test whose p-value threshold scales dynamically with the total number of input LGs, correcting for the increased number of pairwise comparisons in larger datasets.

Connected components of the bipartite graph define candidate ancestral LGs. Where a strict one-to-one correspondence exists between daughter LGs, the ancestral state is inferred directly as the merged content of both vertices. For more complex components indicative of chromosomal fissions or fusions (for example, two LGs in one daughter lineage mapping to a single LG in the other), the pipeline employs outgroup polarisation to resolve the ambiguity. A representative outgroup is dynamically selected from the species tree based on proximity to the node under reconstruction, and its chromosomal configuration is queried to determine the ancestral state via maximum parsimony. Where the outgroup supports a split configuration, the ancestral node is reconstructed as two separate LGs; where it supports a fused configuration, the candidate components are merged. Orphan gene segments that cannot be unambiguously assigned to a primary partition are reassigned based on maximal orthogroup overlap with existing ancestral LGs. The resulting ALG set and its constituent orthogroup membership are saved iteratively and propagate as the reference architecture for all deeper nodes in the reconstruction.

### Automated visualization and output generation

The pipeline includes an automated plotting module built on ggplot2 (Wickham 2016) and RIdeogram (Hao et al. 2020) that generates a standardised set of outputs at every evaluated node. Visual outputs include stacked bar plots of syntenic state distributions across chromosomes, linear karyoplots mapping syntenic blocks onto extant physical chromosomal coordinates, Oxford grid dot plots showing the spatial distribution of orthologs between pairs of species, and ribbon diagrams illustrating the chromosomal rearrangements inferred between lineages. Summary statistics are written to tabular files at each node, reporting the number of inferred ALGs, orthogroup counts per ALG, and the proportion of the gene repertoire assigned to reconstructed linkage groups. These outputs are designed to facilitate both the detection of potential artifacts and the biological interpretation of ancestral architectures across the phylogeny.

### Simulation framework for benchmarking

To quantify the accuracy and sensitivity of LINGUINE across a controlled range of evolutionary scenarios, we developed GARLIC (Genome reARrangement simuLator for Inferring Chromosomal Landscapes), a standalone simulation tool implemented in R. GARLIC evolves a user-defined ancestral genome along a specified species topology, applying stochastic chromosomal rearrangement events (translocations, fissions, and fusions) and gene loss independently along each branch according to user-specified per-branch rates. Each gene in the simulated genome belongs to an orthogroup defined at the ancestral node, allowing the true ALG membership of every gene in every extant species to be recorded and used as ground truth for downstream benchmarking.

For all simulations in this study, the ancestral genome comprised 5,000 orthogroups distributed across 15 chromosomes. Gene copy number per orthogroup was drawn from a shifted geometric distribution (p = 0.9), with gene lengths sampled uniformly between 500 and 2,000 bp. A probability of 0.7 was applied to retain paralogous copies on the same chromosome as their progenitor, reflecting the empirical co-localisation tendency of recent duplicates. All simulations used a three-species topology ((Species A, Species B), Species C). All simulation parameters, including genome size, chromosome number, gene length distributions, copy number probabilities, and per-branch rearrangement rates, are specified in a user-editable YAML configuration file, allowing any parameter to be modified independently to suit the evolutionary context under investigation. For each scenario, the focal evolutionary parameter was varied across its tested range with 100 independent replicates per step, each using a distinct random seed to ensure stochastic independence. Reconstruction accuracy was evaluated by comparing the inferred ALG assignments at Node AB to the known ancestral configuration for all genes in Species A and Species B.

### Comparative performance evaluation

To reconstruct ancestral linkage groups at nematode node N1 using odp, we selected four representative species: *Caenorhabditis elegans* (CELE), *Oscheius tipulae* (OTIP), *Pristionchus exspectatus* (PEXS), and *Strongyloides ratti* (SRAT), chosen to span the diversity of the 13-genome dataset while remaining within the species limit imposed by odp. The minimum scaffold size was set to 4.5 Mb (minscafsize: 4500000). Ancestral LG reconstruction was performed iteratively across all valid three-species combinations from the four selected taxa, always including *C. elegans* as the anchor species. The resulting reciprocal best hit (rbh) files were merged using the *C. elegans* genome as the common reference, and this unified set of ortholog correspondences was used to generate the final pairwise alignments and define the ancestral linkage groups at node N1. The inferred ancestral states were subsequently projected onto the remaining nine species in the 13-genome dataset to annotate linkage groups across the full phylogeny.

Ancestral linkage inference via Syngraph was performed using conserved single-copy markers identified from BUSCO v4.1.4 (Simão et al. 2015) with default parameters using nematoda_odb10 for nematodes and metazoa_odb10 for clitellates) and single copy genes from OrthoFinder v3.1.2 (Emms et al. 2025) with default parameters and as inputs analyses of all 13 nematode genomes. BUSCO results were used to generate a tabular input of orthologous gene coordinates for each species, retaining only genes present as single-copy orthologs across the dataset. The Syngraph infer module was run with *O. tipulae* as the reference taxon (-s OTIP) and a minimum syntenic block threshold of 30 markers (-m 30). The same BUSCO and OrthoFinder-based input files were used for the clitellate annelid dataset, with *N. najaformis* designated as the reference taxon.

LINGUINE was run using HOGs and the rooted species tree produced by OrthoFinder. For both the nematode and clitellate datasets, the minimum scaffold size threshold for ancestral assignment was set to 4.5 Mb. All other parameters were kept at default values as described above. Reconstruction was performed independently for each dataset via full post-order traversal of the respective species tree, with paralogy resolution applied prior to graph construction at each node.

## Data availability

LINGUINE and GARLIC are available in GitHub in the following open repositories, respectively, https://github.com/MetazoaPhylogenomicsLab/Vargas-Chavez-Fernandez_2026_LINGUINE, https://github.com/MetazoaPhylogenomicsLab/Vargas-Chavez-Fernandez_2026_GARLIC

## Author contributions

Carlos Vargas-Chávez: Conceptualization, Methodology, Software, Formal Analysis, Investigation, Data Curation, Visualization, Writing – Original Draft, Writing – Review & Editing.

Rosa Fernández: Conceptualization, Methodology, Visualization, Resources, Funding Acquisition, Project Administration, Supervision, Writing – Original Draft, Writing – Review & Editing.

## Acknowledgements

We thank the members of the Metazoa Phylogenomics and Genome Evolution lab (particularly Dearbhaile Casey) for helpful discussions on the experimental design and performance. R.F. acknowledges support from the European Research Council (grant agreement no. 948281)) and the Agencia Estatal de Investigación (PID2024-161173NB-I00 funded by MICIU/AEI/10.13039/501100011033 and ERDF, EU). The authors declare the usage of AI tools (ChatGPT and Perplexity) to improve text clarity and grammar.

